# Prenatal Alcohol Exposure Impairs Striatal Cholinergic Function and Cognitive Flexibility in Adult Offspring

**DOI:** 10.1101/2025.05.27.656351

**Authors:** William Purvines, Himanshu Gangal, Xueyi Xie, Joseph Ramos, Xuehua Wang, Rajesh Miranda, Jun Wang

## Abstract

Fetal Alcohol Spectrum Disorder (FASD), caused by prenatal alcohol exposure (PAE), is characterized by significant cognitive impairments, including reduced cognitive flexibility. Despite the critical role of cholinergic interneurons (CINs) in the dorsomedial striatum (DMS) for cognitive and behavioral flexibility, their contribution to neurobehavioral deficits in FASD remains unclear. To address this gap, this research explored the impact of PAE on CIN populations and activity, cognitive flexibility, and compulsive drinking behaviors in adult offspring. Using ChAT-Cre;Ai14-tdTomato mice combined with ChAT staining, we found substantial reductions in CIN number within the striatum of adult PAE offspring. Functional assessments revealed that PAE markedly decreased CIN firing activity and reduced acetylcholine (ACh) release in the DMS, as measured by electrophysiology recordings and live-tissue confocal imaging using a genetically encoded ACh sensor. Behaviorally, PAE-exposed offspring exhibited a significant deficit in adapting to reversed action-outcome contingencies despite intact initial learning capabilities. Moreover, PAE-exposed mice exhibited compulsive alcohol drinking behavior, characterized by elevated consumption and preference for quinine-adulterated alcohol. These findings collectively highlight the critical role of impaired cholinergic signaling in the cognitive and behavioral deficits observed following PAE. Understanding this cholinergic dysfunction provides valuable insights necessary for developing targeted interventions aimed at mitigating cognitive and behavioral consequences associated with FASD.

**Highlights:** PAE reduces the number of CINs, their firing rate and ACh release in the DMS.

PAE impaired instrumental reversal learning, known to be mediated by CINs.

PAE increased compulsive-like drinking of quinine adulterated alcohol.

## Introduction

Fetal Alcohol Spectrum Disorder (FASD) encompasses a continuum of neurodevelopmental and neuropsychiatric conditions caused by prenatal alcohol exposure (PAE), affecting an estimated 1.1–9.8% of the U.S. population in school systems (May et al., 2014; May et al., 2018). Individuals with FASD exhibit persistent impairments in cognition, motor function, and brain structure that can extend into adulthood. Among the various cognitive deficits, impaired cognitive flexibility, the capacity to adapt behavior in response to changing environmental contingencies, is a prominent symptom that contributes to impaired executive function (Green et al., 2009; Mattson et al., 1999; McGee et al., 2008). However, despite its clinical salience, the underlying neural mechanisms responsible for this impairment remain poorly understood.

The dorsomedial striatum (DMS) has emerged as a key neural substrate supporting cognitive flexibility (Bradfield et al., 2013; Huang et al., 2024; Ma et al., 2022; Ragozzino et al., 2002). Furthermore, the DMS is vulnerable to the effects of developmental alcohol exposure, with previous studies reporting PAE-induced alterations in striatal plasticity and medium spiny neuron (MSN) function (Cheng et al., 2018; Roselli et al., 2020). Within the DMS, MSNs form the principal output of the striatum, while cholinergic interneurons (CINs), though sparse in number, play a critical regulatory role in shaping striatal circuit dynamics (Gerfen and Surmeier, 2011; Kreitzer, 2009; Kreitzer and Malenka, 2008). CINs are the primary source of acetylcholine (ACh) in the striatum and exert widespread modulatory control over striatal function (Huang et al., 2024; Ma et al., 2022). DMS CINs, whose dynamic firing is shaped by upstream excitatory inputs, are essential for behavioral flexibility, as CIN circuit manipulation alters reversal learning (Bradfield et al., 2013; Li et al., 2025; Ma et al., 2022; Ragozzino et al., 2009). Notably, gestational alcohol exposure has been shown to impair CIN excitability in the dorsolateral striatum (Bariselli et al., 2023). However, the impact of PAE on CINs within the DMS and the behavioral consequences of such alterations remain largely unexplored. Given the established role of DMS CINs in mediating cognitive flexibility (Gerfen and Surmeier, 2011; Kreitzer, 2009; Kreitzer and Malenka, 2008), this is a critical knowledge gap.

Our prior work has demonstrated that alcohol-induced CIN dysfunction in the DMS impairs behavioral flexibility in adult rodents (Ma et al., 2022), raising the possibility that similar mechanisms may underlie cognitive deficits seen following PAE. In this study, we used a preclinical model of FASD by exposing pregnant dams and their pups to prenatal alcohol and found that PAE reduces the population of DMS CINs, suppresses their activity, and decreases ACh release in the DMS in adult offspring. These cellular deficits were accompanied by marked impairments in cognitive flexibility, as evidenced by failure to adapt to reversed action-outcome contingencies, and by increased compulsive drinking of quinine-adulterated alcohol. Our findings suggest that CIN dysfunction in the DMS could be a key neural mechanism underlying behavioral inflexibility and compulsive alcohol-seeking behavior following PAE. These results not only expand our understanding of FASD pathophysiology but also highlight DMS cholinergic circuits as a promising therapeutic target for mitigating cognitive deficits associated with developmental alcohol exposure.

## Materials and Methods

### Animals

Chat-Cre, Ai14, and Ai32 mice were obtained from the Jackson laboratory. Tail DNA samples were collected from mice and PCR was conducted to determine genotype. Mice were housed in same-sex colonies under a 12 h light/dark schedule, with food and water available ad libitum. All animal procedures in this study were approved by the Texas A&M University Institutional Animal Care and Use Committee. All procedures were conducted in agreement with the Guide for the Care and Use of Laboratory Animals, National Research Council, 1996.

### Breeding and Prenatal Alcohol Exposure

For experiments investigating the effect of maternal alcohol drinking on offspring, female mice were trained to consume high levels of alcohol for 6 weeks using the intermittent access to 20% alcohol two-bottle choice (2BC) procedure and were then mated with alcohol-naïve male mice as previously described (Cheng et al., 2018; Gangal et al., 2023; Huang et al., 2024; Iannucci et al., 2023; Ma et al., 2022; Wang et al., 2023; Xie et al., 2023a; Xie et al., 2023b). In this procedure, mice were given 24-hr access to 1 bottle of 20% alcohol in water (vol/vol) and 1 bottle of water every other weekday (Monday, Wednesday, Friday). Alcohol solutions were prepared by diluting 200-proof pure ethanol in drinking water. Bottle placement was altered each session to control for side preference. Using this protocol, we have previously observed that pregnant damns reach alcohol intake levels of 20-30 g/kg over 24 hours (Cheng et al., 2018). For the control (Ctrl) group, female mice were given free access to water only. During mating, only water was administered to prevent males from drinking. After mating, impregnated females were re-exposed to alcohol using the 2BC procedure until postnatal day 10. PAE and Ctrl offspring were separated by sex at 21 days old and housed in groups of 3-5.

In experiments conducted for figures 2 and 3, maternal alcohol vapor inhalation was used to expose pups to prenatal alcohol. This exposure paradigm was utilized as alcohol vapor inhalation is known to achieve higher, more reliable blood alcohol concentrations (BACs) compared to voluntary drinking models which often produce lower, more variable BACs (Gilpin et al., 2008). Pregnant dams were either exposed to air or alcohol vapor pumped into a plexiglass chamber (92 x 62 x 36 cm) for 1.5-2 hours on embryonic days 11-15 to produce Ctrl and PAE offspring, respectively. Our vapor inhalation set up was adapted from (Morton et al., 2014) and has been previously described (Vierkant et al., 2023b). To produce ethanol vapor, air was pumped into a sealed flask containing 200 proof alcohol via an aeration stone, which was then mixed with air before entering the chamber. In trial experiments, this vapor exposure induced BACs of ∼120 mg/dL. After pups were born, they were separated and housed as described in the two-bottle choice procedure.

**Figure 1.**
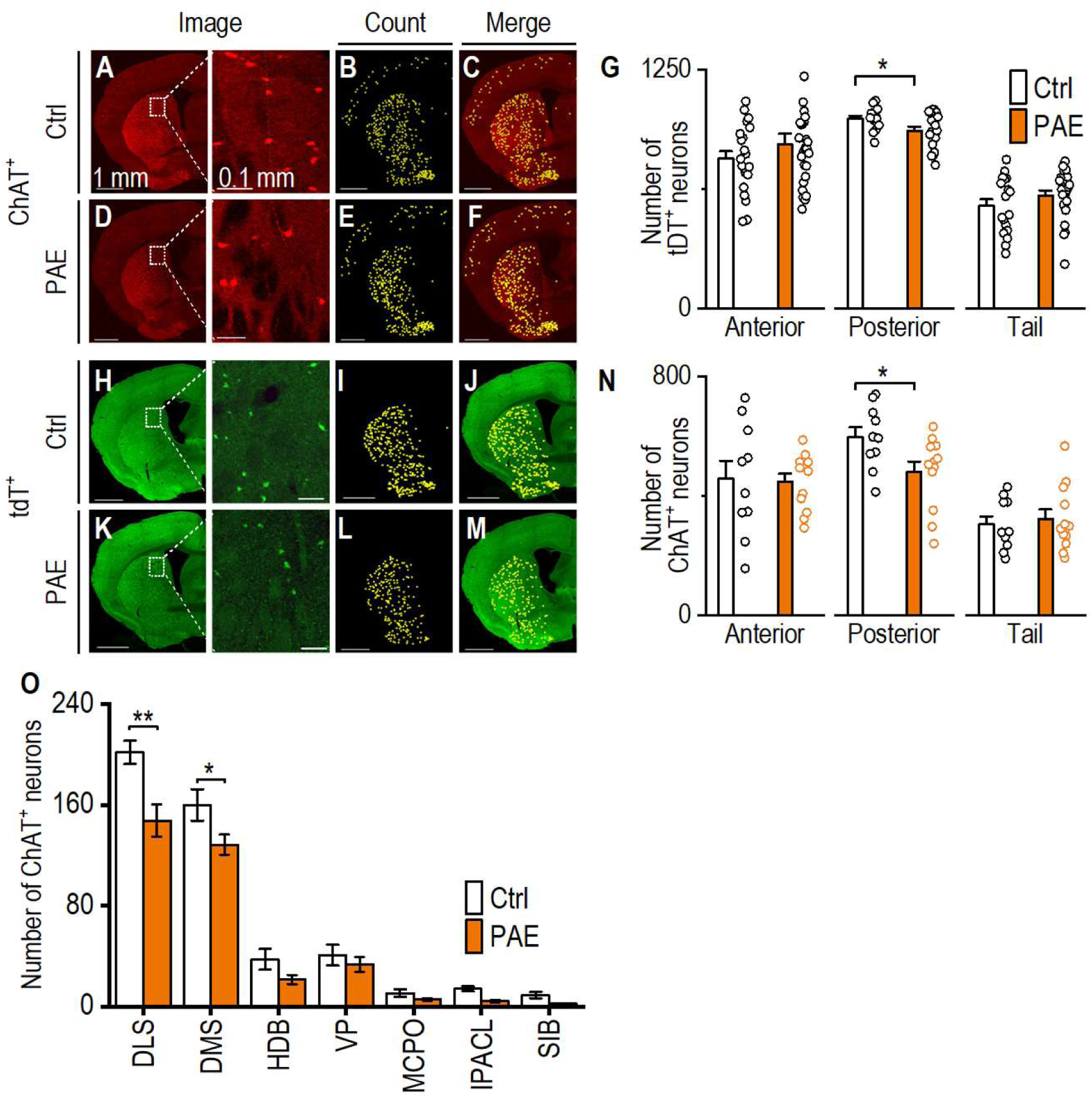
Prenatal alcohol exposure reduces cholinergic neurons in adult offspring. ChAT-Cre;Ai14 mice were prenatally exposed to alcohol, and coronal brain sections were analyzed at 8 months of age. Selected sections were also stained with anti-ChAT antibody. ***A****-**F***, Sample images showing tdTomato (tdT)^+^ neurons (*A*, *D*), spots of counted neurons (*B*, *E*), and merged images of neurons and spots (*C*, *F*) in a posterior section (0.74 mm relative to bregma) from control (Ctrl, *A*-*C*) and prenatal alcohol exposed (PAE, *D*-*F*) groups of mice. Enlarged images in *A* and *D* show fewer tdT^+^ neurons in the dorsomedial striatum (DMS) of a PAE mouse (*D*) than in that of a Ctrl mouse (*A*). Each yellow spot in *B* and *E* represents a tdT^+^ neuron in *A* and *D*, respectively. ***G***, Bar graph comparing tdT^+^ neuron counts in anterior (1.70 to 0.62 mm), posterior (0.50 to −0.46 mm), and tail (−0.58 to -2.06 mm) brain sections. \**p* < 0.05; n.s., not significant (p > 0.05) by un-paired *t* test. n = 21 sections from 3 mice (21/3) (Anterior Ctrl); 27/4 (Anterior PAE); 14/3 (Posterior Ctrl), 19/4 (Posterior PAE), 18/3 (Tail Ctrl), and 24/4 (Tail PAE). ***H***-***M***, Representative images of ChAT-immunoreactive (ChAT^+^) neurons (*H*, *K*), spots of counted ChAT^+^ neurons (*I*, *L*), and merged images (*J*, *M*) in posterior sections from Ctrl (*H*-*J*) and PAE (*K*-*M*) mice. Note that enlarged images in *H* and *K* show fewer ChAT^+^ neurons in the DMS of a PAE mouse (*K*) than in that of a Ctrl mouse (*H*). ***N***, Bar graph summarizing ChAT^+^ neuron counts in anterior, posterior, and tail sections for the control and PAE groups. \**p* < 0.05 by un-paired *t* test. n = 10/3 (Anterior Ctrl); 12/4 (Anterior PAE); 10/3 (Posterior Ctrl); 12/4 (Posterior PAE); 10/3 (Tail Ctrl); 12/4 (Tail PAE). The 3 Ctrl mice were derived from 3 litters and the 4 PAE mice were derived from 4 litters. ***O***, Bar graph comparison of tdT^+^ neurons between Ctrl and PAE mice across multiple brain regions. \**p* < 0.05, \*\**p* < 0.01 by unpaired *t* test, n = 10/5, Ctrl; 10/5, PAE. The 5 Ctrl mice were derived from 3 litters and the 5 PAE mice were derived from 5 litters. DLS, dorsolateral striatum; VP, ventral pallidum; HDB, horizontal diagonal band; IPACL, interstitial nucleus of the posterior limb of the anterior commissure lateral part; MPCO, magnocellular preoptic nucleus; SIB, Substantia innominata-basal part.

**Figure 2.**
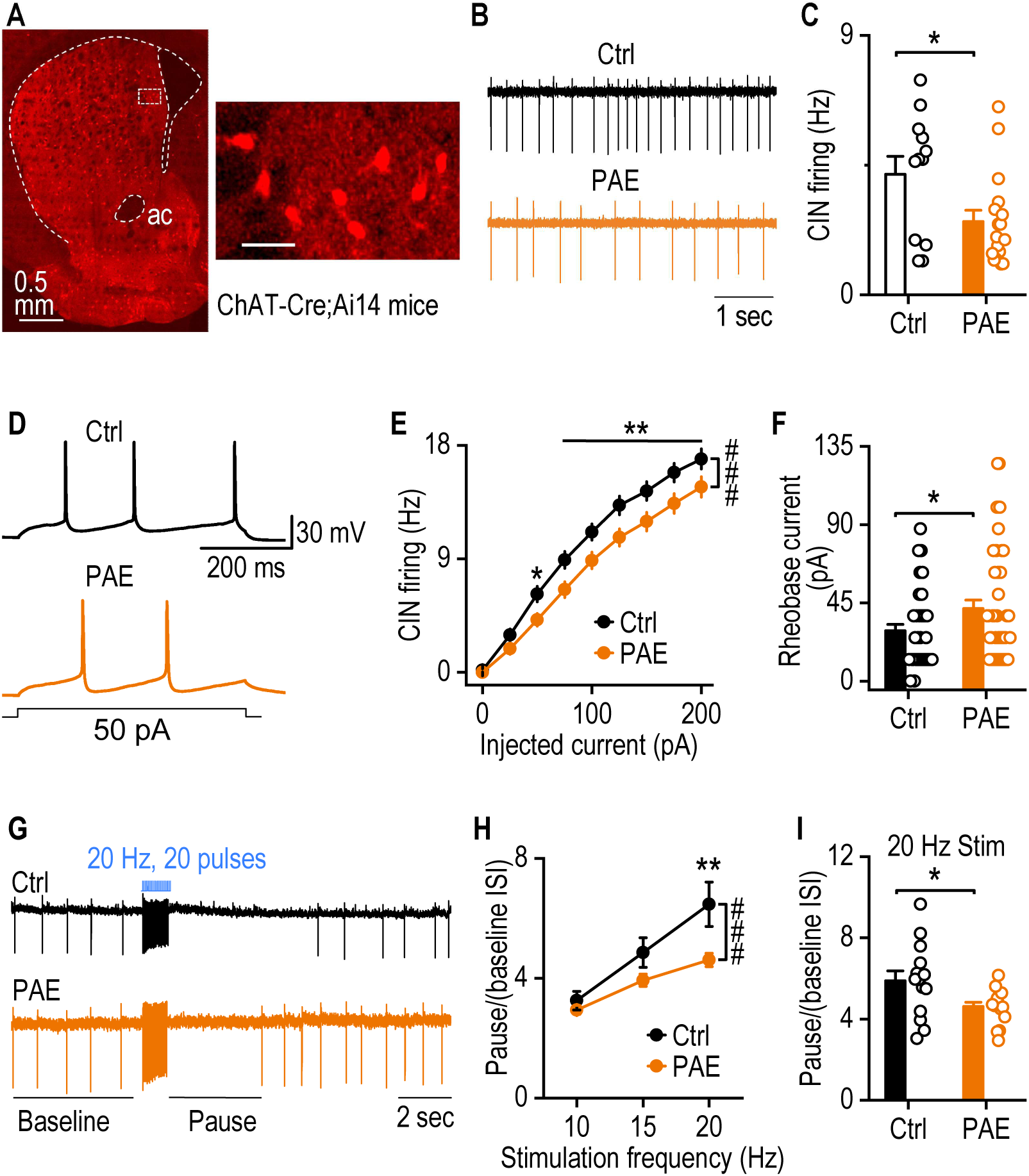
PAE reduces CIN firing activity in the DMS of adult offspring. ***A***, Representative image of a ChAT-Cre;Ai14 mouse showing tdT expression in CINs of the DMS. ac, anterior commissure. Scale bar: 50 µm (inset). ChAT-Cre;Ai14 mice were exposed to alcohol both prenatally and for 10 days postnatally. DMS slices were collected at 8 months of age. ***B***, Representative traces of spontaneous CIN firing in Ctrl and PAE groups recorded using cell-attached configuration. ***C***, PAE significantly decreases the spontaneous firing rate of DMS CINs in adult offspring. \**p* < 0.05, unpaired t-test. n = 12 neurons from 3 mice (12/3) that were derived from 3 litters for Ctrl and 17/3, derived from 3 litters, for PAE. ***D***, Sample traces of evoked CIN firing in Ctrl and PAE groups. ChATCre;Ai32 mice were exposed to air or alcohol vapor prenatally. ***E***, PAE significantly reduced evoked firing of DMS CINs in response step-current injection. ^###^*p* < 0.001; \**p* < 0.05, \*\**p* < 0.01 versus PAE at the same stimulating intensities, by 2-Way RM ANOVA with post-hoc Tukey tests. n= 37 neurons from 4 mice (37/4) for Ctrl and 46/4 for PAE. ***F***, PAE mice exhibited higher rheobase currents than Ctrl animals. Rheobase current was defined as the first current step, within a series of +25 pA steps beginning at 0 pA, capable of eliciting one action potential. \**p* < 0.05 by unpaired t test n= 37/4 (Ctrl) and 46/4 (PAE). ***G***, Representative traces from a Ctrl and PAE mouse showing burst-pause responses in CINs evoked by optogenetic stimulation. ***H***, PAE significantly reduced normalized pause durations compared to Ctrl mice. The pauses were induced by escalating frequencies of optogenetic stimulation (each lasting 1 sec). The normalized pause duration was calculated by dividing the raw pause duration by the average baseline inter-spike interval (ISI) prior to stimulation. \*\**p* < 0.01, ^###^*p* < 0.001, Group × Stimulation interaction by two-way repeated measures ANOVA with Tukey’s post-hoc tests. n = 14 neurons from 4 mice (14/4) for Ctrl and 15/4 for PAE. ***I***, Comparison of normalized pause duration following 20 Hz stimulation between Ctrl and PAE mice. CINs from PAE mice exhibited significantly shorter normalized pause durations than those from Ctrl mice. \**p* < 0.05, unpaired t-test. n = 14 neurons from 4 mice (14/4) for Ctrl and 15/4 for PAE.

### Histology

Mice were anesthetized and perfused with 4% paraformaldehyde (PFA) in phosphate-buffered saline (PBS). Brains were collected and submerged in 4% PFA/PBS solution overnight at 4°C before being transferred to 30% sucrose in PBS. After brains sunk, they were sliced on a cryostat into 50 µm sections. Some sections were stained with anti-ChAT antibody (AB144P) followed by a secondary antibody conjugated to a 647nm-emmiting fluorophore (A21447, Alexa-Fluor 647) (Gangal et al., 2023). Confocal images were obtained with an Olympus Fluoview 1200 microscope using a 10x objective lens. Images were stitched with the same program and fluorescent neurons were counted using Imaris.

### Electrophysiology

Electrophysiological recordings were conducted as previously described (Gangal et al., 2023; Huang et al., 2024; Ma et al., 2022). Mice were perfused, and 250 μM DMS-containing slices were collected in ice-cold cutting solution. Cutting solution contained (in mM): 40 NaCl, 148.5 sucrose, 4 KCl, 1.25 NaH^2^PO_4_, 25 NaHCO^3^, 0.5 CaCl_2_, 7 MgCl_2_, 10 glucose, 1 sodium ascorbate, 3 sodium pyruvate, and 3 myoinositol, saturated with 95% O_2_ and 5% CO_2_. Slices were incubated in a 1:1 mixture of cutting and external solution held at 32°C for 45 minutes before being transferred to pure external solution at room temperature for 15 minutes before use and the duration of the experiment. External solution consisted of (in mM): 125 NaCl, 4.5 KCl, 2.5 CaCl_2_, 1.3 MgCl_2_, 1.25 NaH_2_PO_4_, 25 NaHCO_3_, 15 sucrose, and 15 glucose, saturated with 95% O_2_ and 5% CO_2_.

In the recording chamber, slices were perfused with 32°C external solution at a flow rate of 1.5-2 mL/min. CINs were identified by their Cre-driven expression of fluorescent reporter and their large soma size. All recordings were conducted using a Multiclamp 700B amplifier controlled by pClamp 11.4 software (Molecular Devices). In both whole-cell patch clamp and cell-attached recordings, a K^+^ intracellular solution was used, consisting of (in mM): 123 potassium gluconate, 10 HEPES, 0.2 EGTA, 8 NaCl, 2 MgATP, 0.3 NaGTP (pH 7.3), with an osmolarity of 270–280 mOsm.

To measure excitability in whole-cell patch clamp recordings, CINs were recorded in current-clamp mode and action potentials were evoked with stepped current injections at 25-pA increments lasting 500ms. CIN spontaneous and burst-pause firing was measured using cell-attached recording in voltage clamp mode. For optogenetic excitation in burst-pause recordings, 10, 15, and 20 Hz blue light stimulation (473 nm, 2 ms pulse width) was delivered through the objective lens.

### Live tissue confocal imaging of ACh release

A green ACh sensor, AAV-gACh4m (Huang et al., 2024; Jing et al., 2020), was bilaterally infused into the DMS of ∼4 month old WT PAE or Ctrl mice (AP: 0.1, ML: ±1.87, DV: -2.9mm). 2 weeks later, acute brain slices were collected in the same manner as in electrophysiology experiments. Slices were held in a recording bath continuously perfused with ACSF saturated with 95% O_2_ and 5% CO_2_. ACh release was imaged with an Olympus Fluoview FV3000 microscope using a 40x NA 0.8 water immersion objective along with a 488 nm and 561 nm laser. A sample rate of ∼2 frames per second was used for imaging, and all imaging parameters were kept consistent throughout all imaging sessions. MATLAB scripts were used to extract time-course ΔF/F values to quantify spontaneous event frequency and release amplitude.

### Instrumental reversal learning

The instrumental reversal learning procedure was conducted as described previously (Gangal et al., 2023; Huang et al., 2024; Ma et al., 2022). Training began with either a purified pellet or grain pellet sub-session. During these sessions, the house light was on, and one of the levers was presented. Initially, pressing the left lever (A1) resulted in delivery of a grain pellet reward (O1), while pressing the right lever (A2) delivered a purified pellet reward (O2). Animals were first trained under a fixed ratio 1 (FR1) schedule for three days, where each lever press yielded one reward. The mice were then transitioned to a random ratio (RR) schedule, beginning with RR5 (probability of reward = 0.2 per press) for three days, then progressing to RR10 for another three days, and finally progressing to RR20 for five days. The initial devaluation test occurred one day after the final RR20 training session. Prior to the test, animals had one-hour free access to 40 pellets (randomly assigned as either purified or grain) to induce satiety. The test began either 10 minutes after consuming all the pellets or after a maximum of 60 minutes. During the 10-minute session, both levers were presented, but lever presses did not result in reward delivery. The lever corresponding to the pre-fed pellet type was classified as devalued (Dev), while the lever associated with the pellet type not pre-fed was classified as valued (Val). The test procedure was repeated with the alternative pellet type on the following day.

PAE and Ctrl mice were then exposed to four sessions of reversal learning. During reversal, action-outcome contingencies were switched: the left lever (initially A1→O1 paired with grain pellet O1) now delivered purified pellet reward (A1→O2), and the right lever (initially A2→O2 paired with purified pellet O2) delivered grain pellet reward (O1). Another devaluation test was then conducted to asses how well mice learned the reversed contingencies. This second devaluation test followed the same procedure as the initial devaluation test. The devaluation index was calculated as (Val - Dev) / (Val + Dev).

### Intermittent Access Two-bottle Choice Drinking Procedure

Two-bottle choice drinking was conducted as previously described (Cheng et al., 2017; Cheng et al., 2018; Gangal et al., 2023; Ma et al., 2022; Vierkant et al., 2023a; Xie et al., 2023b). Adult PAE and Ctrl mice were individually housed to assess alcohol drinking. Mice were presented with two bottles: one containing a 20% alcohol solution in drinking water and the other containing drinking water only, provided three times a week (Monday, Wednesday, Friday) for 24 hours. Both sets of bottles were replaced with fresh solutions weekly. To measure alcohol consumption, bottles were weighed before and after each session, and alcohol intake was calculated as the grams of alcohol consumed divided by the mouse’s body weight in kilograms (g/kg/24h). Alcohol preference was determined as the total alcohol solution intake divided by the total fluid intake, multiplied by 100%. To account for drippage, additional water and alcohol bottles were placed over empty cages during sessions. In experiments examining compulsive drinking behavior, quinine hemisulfate salt was added to the 20% alcohol solution at a concentration of 50 mg/L.

### Statistical Analysis

Data from male and female subjects were combined for analysis and sex differences were not evaluated. All data were analyzed using unpaired *t* tests and two-way ANOVA with repeated measures (two-way RM ANOVA), followed by Tukey’s *post-hoc test*. Statistical analysis was conducted by SigmaPlot. All data were expressed as the Mean ± SEM.

## Results

### PAE Reduces Cholinergic Neuron Populations Across Multiple Brain Regions of Adult Offspring

To determine the impact of PAE on the number of cholinergic neurons, pregnant Ai14 (Cre-dependent tdTomato) dams were exposed to 2-bottle choice alcohol or water and crossed with male ChAT-Cre sires to produce PAE or Ctrl ChAT-Cre;Ai14 offspring, respectively (Cheng et al., 2018). Mice from both groups were perfused at ∼8 months of age. Coronal sections were prepared and scanned with a confocal microscope and the number of tdTomato-positive (tdT^+^) neurons were counted. We examined sections from 3 different anterior-posterior planes: anterior (Ant., AP: 0.90 ± 0.04), posterior (Post., AP: 0.09 ± 0.06), and tail (Tail, AP: -1.13 ± 0.05). We found that the number of tdT^+^ neurons at the posterior plane was significantly lower in the PAE group than in Ctrl subjects (Fig. 1A-1G; *t*_(31)_ = 2.21, *p* < 0.05). However, we did not find any difference in the number of tdT^+^ neurons at anterior and tail planes between Ctrl and PAE groups (Fig. 1G; Ant.: *t*^(46)^ = -0.71, *p* > 0.05; Tail: *t*_(40)_ = -1.28, *p* > 0.05) tdT^+^ neurons in ChAT-Cre;Ai14 mice represent neurons with a history of ChAT expression, and some of these neurons may lose ChAT expression as the offspring age. To further confirm the PAE-mediated reduction of the number of cholinergic neurons, we also stained the sections from these mice with an anti-ChAT antibody and then imaged and counted these ChAT-immunoreactive (ChAT^+^) neurons. Again, we found that the number of ChAT^+^ neurons at the posterior plane was significantly lower in the PAE group than their water controls (Fig. 1H-1N; *t*_(20)_ = 2.38, *p* < 0.05). Immunoreactive neuron numbers in the anterior and tail planes did not differ between control and PAE groups (Fig. 1N; Ant.: *t*_(20)_ = 0.15, *p* > 0.05; Tail: *t*_(20)_ = -0.44, *p* > 0.05). We then analyzed ChAT^+^ neuron expression in specific regions of the posterior plane and found that cholinergic neuron numbers in the dorsomedial striatum and dorsolateral striatum were significantly lower in the PAE than control mice (Fig. 1O, dorsolateral striatum: *t*_(8)_ = 3.42, *p* < 0.01 ; dorsomedial striatum: *t*_(8)_ = 2.14, *p* < 0.05). These results indicate that PAE significantly reduces the number of cholinergic neurons at the posterior plane of adult offspring.

### PAE Reduces CIN Firing Activity in the DMS of Adult Offspring

We next investigated whether PAE affects the function of the remaining CINs in the DMS. To address this, ChAT-Cre;Ai14 mice were utilized so that CINs could be easily identified. Dams were prenatally exposed to alcohol or water using the intermittent-access two-bottle choice drinking procedure previously described. DMS slices were prepared from ∼8-month-old offspring, and CINs were identified by their Cre-driven expression of tdTomato (Fig. 2A). Using cell-attached recording, we found that the spontaneous firing frequency of CINs was significantly reduced in PAE mice compared to the Ctrl group (Fig. ^2^B, 2C; *t*_(27)_ = 2.35, *p* < 0.05), indicating that PAE diminishes CIN firing activity in the DMS of adult offspring.

Using our vapor model of alcohol exposure, we next examined whether PAE affects the intrinsic excitability of DMS CINs in ChAT-Cre;Ai32 mice. Whole-cell patch-clamp recordings were performed on DMS slices from ∼5-month-old Ctrl and PAE offspring. A series of 500-ms step currents were injected via the recording electrode to evoke action potentials in CINs (Fig. 2D). We observed that the same current injection elicited a lower firing frequency in the PAE group compared to the Ctrl group (Fig. 2E; *F*_(1,81)_ = 6.54, *p* < 0.05). Consistent with this reduced excitability, the rheobase current was significantly higher in PAE mice than in Ctrl subjects (Fig. 2F; *t*_(81)_ = -2.04, \**p* < 0.05), indicating greater synaptic input is required to excite CINs in PAE offspring.

CINs exhibit a characteristic burst-pause firing pattern in response to excitatory input. To examine whether PAE affects this dynamic firing pattern, we recorded CIN spontaneous activity in cell-attached mode before, during, and after inducing burst firing with optogenetic stimulation using a series of blue-light pulses. We found that the pause duration, normalized to the baseline inter-spike interval, was significantly reduced in CINs from PAE mice compared to the Ctrl group. (Fig. 2G, 2H: group x simulation, *F*_(2,54)_ = 8.49, ^#^*p* < 0.001; 2I: *t*_(7)_ = 2.41, \**p* < 0.05). Collectively, these findings suggest that PAE impairs CIN spontaneous and evoked firing in the DMS as well as their characteristic pause in response to burst stimulation.

### PAE Reduces ACh Release in the DMS of Adult Offspring

Given our findings of disrupted CIN physiology following PAE, and considering CINs are the primary source of striatal ACh, we next examined whether PAE alters ACh release in the DMS. A green ACh biosensor, gACh4M, was infused into the DMS of mice exposed to prenatal air (Ctrl) or alcohol vapor (PAE) (Fig. Figure 3A). We then conducted live confocal imaging two weeks later. We found that the amplitude of spontaneous ACh release events was significantly lower in PAE mice compared to Ctrl animals (Fig. Figure 3B, Figure 3C; *t*_(4)_ = 2.18, *p* < 0.05). However, the frequency of spontaneous ACh release events was not significantly affected by PAE (Fig. Figure 3B, Figure 3D; *t*_(14)_ = -0.58, *p* > 0.05). These findings suggest that PAE impairs CIN-driven release of ACh in the DMS.

**Figure 3.**
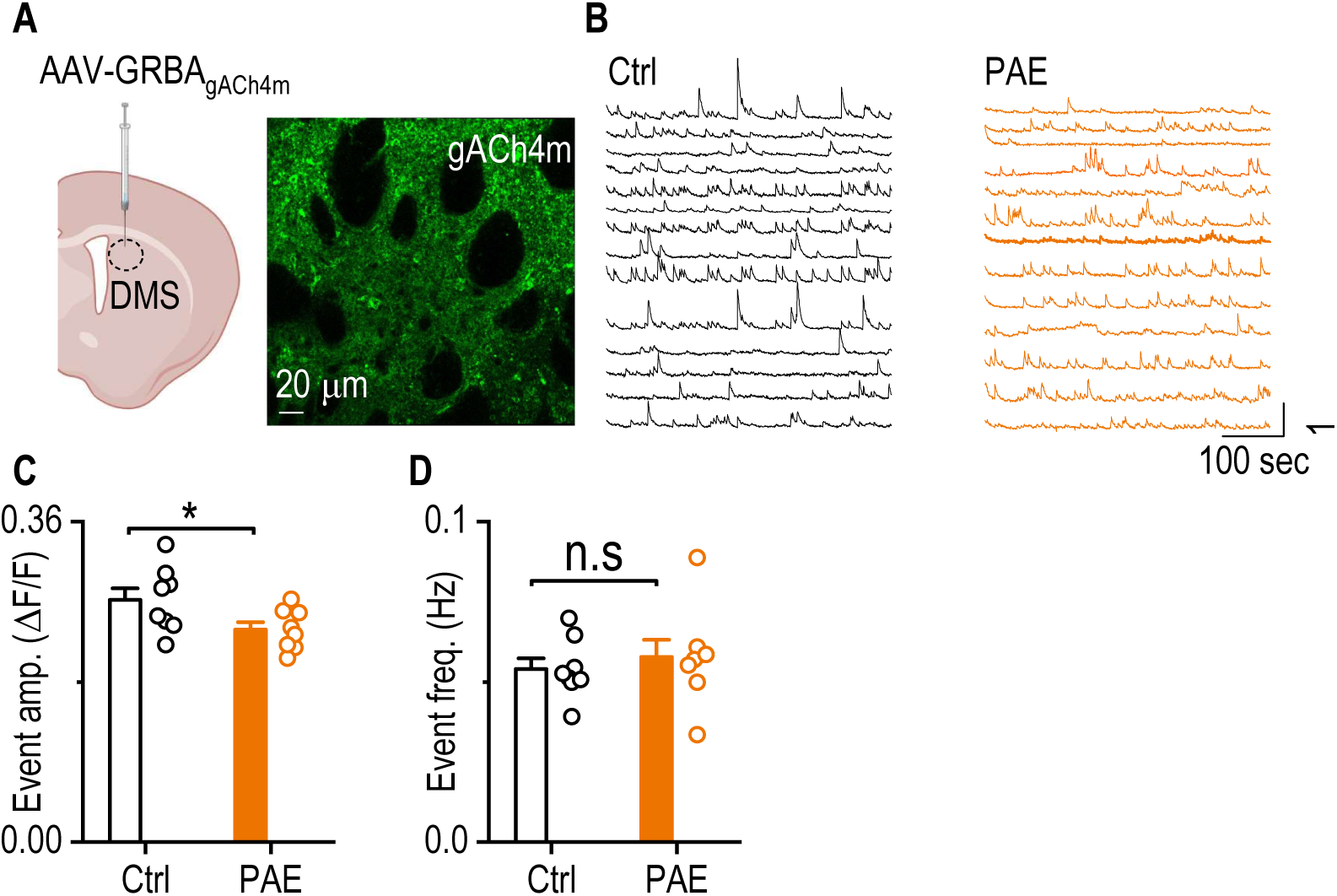
PAE reduces ACh release in the DMS of adult offspring. ***A***, Schematic illustrating viral infusion of AAV-gACh4m into the DMS (left) and a representative image showing the expression of the ACh sensor GRAB-ACh4M (right). Pregnant C57BL/6 dams were exposed to alcohol vapor or air for 5 days during embryonic days 11–15. At ∼4 months of age, offspring from each group received DMS infusions of AAV-gACh4m. DMS slices were prepared 2 weeks later for live confocal imaging. ***B***, Representative traces of ACh release events recorded via live-tissue confocal imaging in Ctrl and PAE groups. ***C, D***, PAE significantly reduced the amplitude (C) but not the frequency (D) of ACh release events in DMS slices from adult offspring. \**p* < 0.05, unpaired t-test. n = 8 mice from 3 litters (Ctrl) and 8 mice from 2 litters (PAE).

### PAE Impairs Cognitive Flexibility in Reversal Learning Tasks of Adult Offspring

Previous research demonstrates that DMS CIN activity is integral for behavioral flexibility (Bradfield et al., 2013; Ragozzino et al., 2009), and it is well known that individuals with FASD display persistent cognitive flexibility deficits into adulthood (Green et al., 2009; Mattson et al., 1999; McGee et al., 2008). Given our findings of impaired cholinergic function in PAE offspring, we investigated whether PAE impairs cognitive flexibility in adult offspring using an operant reversal learning paradigm. Mice were first trained to acquire initial action-outcome (A-O) contingencies, with left lever presses delivering purified pellets to one magazine and right lever presses delivering grain pellets to another. Acquisition of this initial A-O contingency was confirmed by an outcome devaluation test (Fig. Figure 4A). During reversal training, reward contingencies were reversed, such that left-port lever presses delivered grain pellets, and right-port lever presses delivered purified pellets. Another devaluation test was conducted in the same manner following four days of reversal training to evaluate if subjects learned the new A-O contingencies. (Fig. Figure 4B).

**Figure 4.**
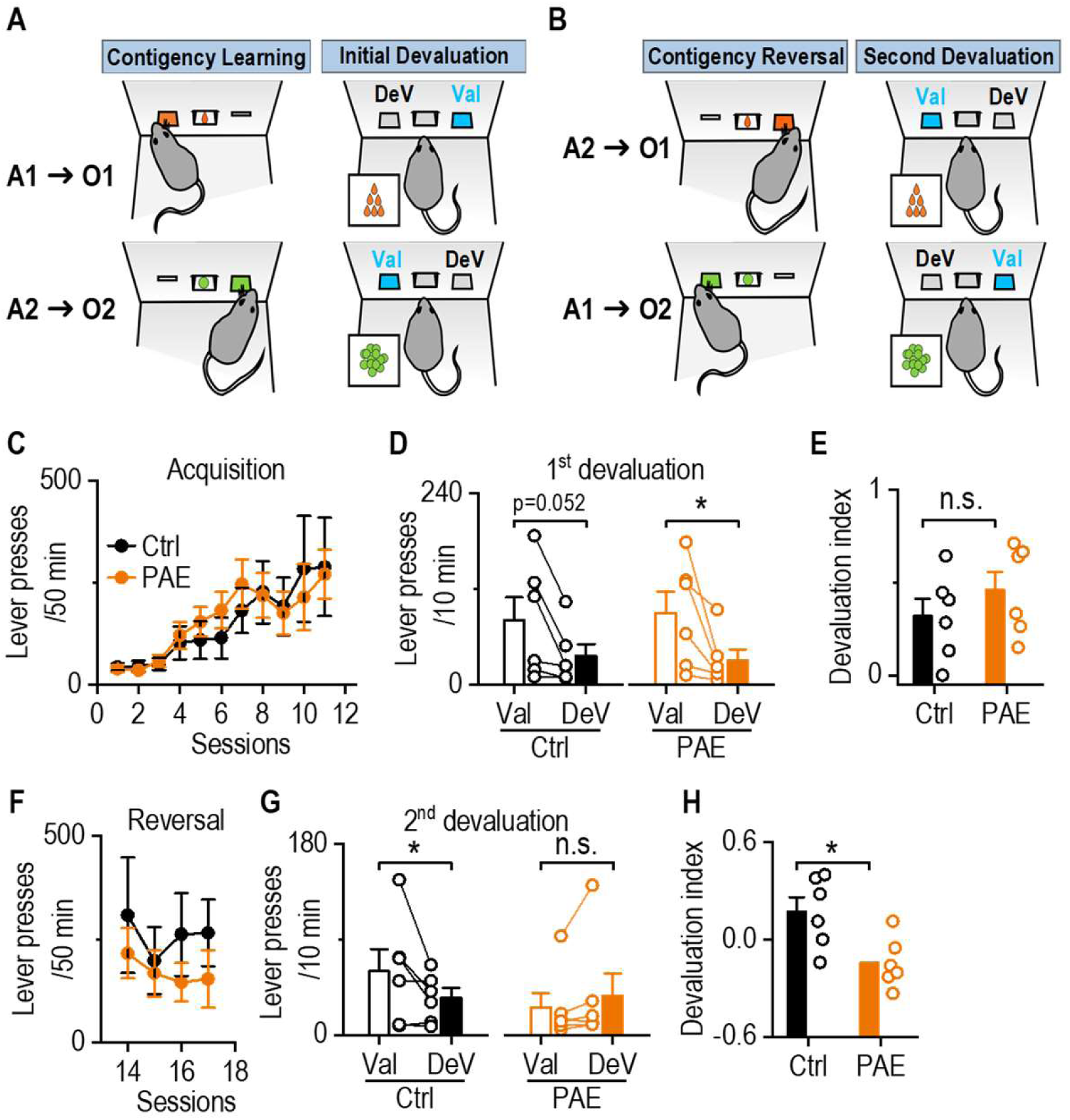
PAE impairs reversal learning of instrumental conditioning in adult offspring. ***A***, Schematic of initial action-outcome (A-O) contingency learning in an operant task. Mice underwent 11 training sessions to press the left lever (A1) for a grain pellet reward (O1) and the right lever (A2) for a purified pellet reward (O2). A devaluation test followed, where each reward was pre-fed before an operant test (devalued) to assess A-O contingency learning. Val, valued; Dev, devalued. ***B.*** Schematic of reversal learning, where A1 was now paired with O2 and A2 with O1. A second devaluation test assessed the learning of reversed A-O contingencies. ***C***, Initial training lever press rates were similar between PAE and Ctrl mice. n.s., not significant p > 0.05 by 2-Way RM ANOVA,. ***D***, Both groups were sensitive to outcome devaluation following initial training. \**p* < 0.05, 2-way RM ANOVA. ***E***, During the first devaluation test, PAE did not affect the devaluation index, calculated as (Val– Dev)/(Val+Dev). n.s., not significant, *p* > 0.05, unpaired t-test. ***F***, Lever pressing rates during reversal learning training were comparable between groups. ***G***, PAE mice exhibited reduced sensitivity to outcome devaluation after reversal learning compared to Ctrl mice. \**p* < 0.05, 2-way RM ANOVA. ***H***, The devaluation index was significantly lower in PAE mice compared to Ctrl mice during the second devaluation test. \**p* < 0.05, unpaired t-test, n = 6 mice from 4 litters (Ctrl) and 6 mice from 4 litters (PAE)

Both groups had similar lever-press rates during initial training, with no significant interaction between group and session (Fig. Figure 4C; *F*_(11,110)_ = 0.38, *p* > 0.05). PAE did not affect the formation of the initial A-O contingencies, as PAE and Ctrl mice showed comparable sensitivity to devaluation in the first devaluation test (Fig. Figure 4D; *F*_(1,10)_ = 0.24, *p* > 0.05). Furthermore, the devaluation index did not differ between Ctrl and PAE mice during this first test (Fig. Figure 4D*; t*_(10)_ = -1.04, *p* > 0.05). However, after four sessions of reversal training (Fig. Figure 4F), the PAE group was significantly less sensitive to the second devaluation compared to Ctrl subjects. Ctrl mice significantly reduced their pressing for the devalued lever, while PAE mice pressed similarly for both the valued and devalued levers (Fig. Figure 4G; *F*_(1,10)_ = 5.76, *p* < 0.05). Consistent with this finding, the devaluation index of the PAE group was significantly lower than that of the Ctrl group during this second devaluation test (Fig. Figure 4H; *t*_(10)_ = 2.89, *p* < 0.05). These findings suggest that PAE impairs instrumental behavioral flexibility in adult offspring, as evidenced by reduced sensitivity to the second devaluation following reversal learning. The selective deficit observed after reversal learning indicates that PAE-associated cognitive impairments are specifically related to cognitive flexibility, rather than generalized learning deficits.

### PAE Promotes Compulsive Alcohol Consumption in Adult Offspring

Having found that PAE offspring exhibit deficits in reversal learning, we next investigated whether PAE mice show other relevant deficits in behavioral flexibility. A cardinal feature of alcohol use disorder (AUD) in humans is compulsive consumption and seeking of alcohol (Koob and Volkow, 2010). Furthermore, those with FASD show increased susceptibility to developing substance use disorders (Baer et al., 1998; Goldschmidt et al., 2019). Thus, we investigated whether PAE affected voluntary alcohol drinking in adult offspring. To this end, PAE and Ctrl ChAT-Cre;Ai14 mice underwent an intermittent-access to two-bottle choice procedure with 20% alcohol, a frequently used model of AUD. Surprisingly, we found no significant differences between Ctrl and PAE mice in alcohol intake (Fig. 5A; *F*_(1,13)_ = 2.89, *p* > 0.05) or preference (Fig. 5B; *F*_(1,13)_ = 0.47, *p* > 0.05).

**Figure 5.**
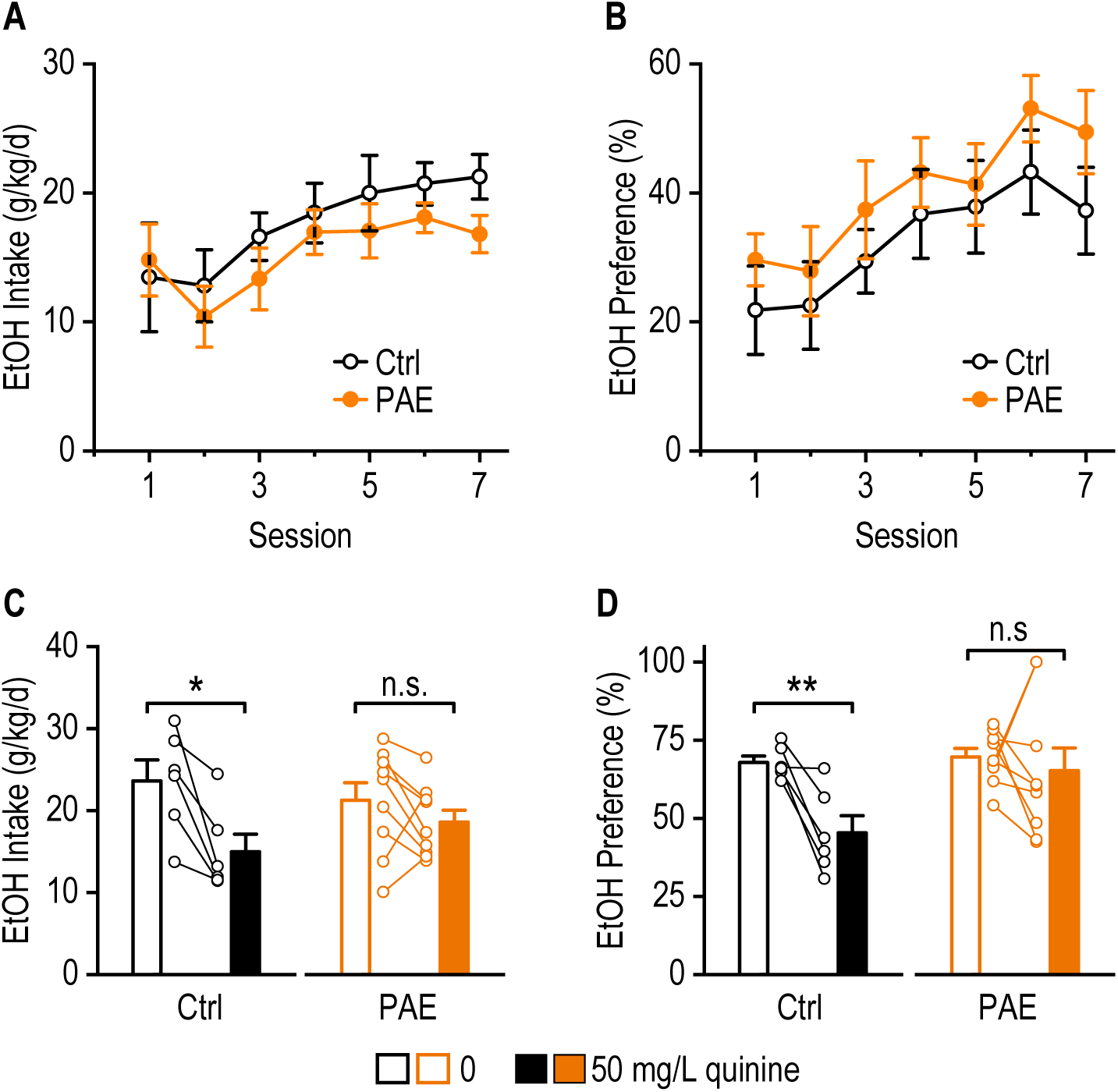
PAE reduces sensitivity to quinine-adulterated alcohol. PAE and Ctrl ChATCre;Ai14 mice (6 months old) were trained to consume alcohol using an intermittent-access two-bottle choice procedure with 20% alcohol. ***A***, PAE did not affect alcohol consumption. *p* > 0.05 by 2-way RM ANOVA ***B***, PAE had no effect on preference for 20% alcohol. *p* > 0.05 by 2-way RM ANOVA. ***C***, Ctrl, but not PAE mice, significantly reduced alcohol intake when 50 mg/L quinine was added. \**p* < 0.05 by unpaired t-test. ***D***, Preference for alcohol was significantly decreased by 50 mg/L quinine in Ctrl mice but remained unchanged in PAE mice. \**p* < 0.05 by unpaired t-test. n = 6 mice from 4 litters (Ctrl) and 9 mice from 5 litters (PAE).

We next hypothesized that PAE mice may specifically show inflexible, compulsive alcohol drinking behavior. To evaluate this, we adulterated the alcohol solution with 50 mg/L quinine, a bitter compound typically causing aversion and reduced alcohol intake. Following quinine adulteration, Ctrl mice reduced alcohol intake when quinine was added, but PAE mice were insensitive to the addition of quinine and did not significantly reduce their drinking (Fig. 5C; Ctrl: *t*_(5)_ = 3.72, *p* < 0.05; PAE: *t*_(8)_ = 1.42, *p* > 0.05). Consistent with this finding, alcohol preference was significantly reduced in Ctrl mice but remained unchanged in PAE mice after quinine addition (Fig. 5D; Ctrl: *t*_(5)_ = 4.40, *p* < 0.01; PAE: *t*_(8)_ = 0.55, *p* > 0.05). Together, these findings indicate that PAE exposure leads to increased compulsive alcohol drinking, suggesting diminished sensitivity to negative outcomes.

## Discussion

The present study provides compelling evidence that PAE induces significant and enduring impairments in CIN populations and cholinergic function in the DMS, associated with deficits in cognitive flexibility and increased compulsive alcohol-drinking behavior in adult offspring. Specifically, we observed marked reductions in the number of striatal CINs, diminished spontaneous and evoked firing rates of CINs, reduced ACh release, impaired performance in reversal learning tasks, and increased compulsive drinking behavior in PAE adult offspring. These findings identify disrupted cholinergic signaling in the DMS as a key neurobiological substrate potentially underlying behavioral inflexibility and compulsivity observed in FASD.

A critical finding of this study is the significant reduction in CIN populations within the striatum following PAE. Although CINs represent a small proportion of striatal neurons, their profound modulatory influence over MSNs and their essential role in shaping striatal network dynamics makes this loss functionally significant. Our results align with previous studies showing vulnerability of striatal circuits, particularly cholinergic neurons, to developmental alcohol exposure (Smiley et al., 2021).

Beyond reductions in neuronal number, our study also demonstrates that the functional integrity of surviving CINs is compromised following PAE. Electrophysiology experiments revealed significant reductions in spontaneous CIN firing rate and diminished intrinsic excitability, indicating that CINs in PAE animals require stronger synaptic input to generate firing. Interestingly, this is consistent with previous work from our laboratory demonstrating that adult exposure to alcohol and cocaine also reduced CIN spontaneous firing (Gangal et al., 2023). Additionally, our findings extend previous observations of CIN excitability deficits reported in the dorsolateral striatum following gestational alcohol exposure (Bariselli et al., 2023). The shortened CIN pause response following excitation is consistent with altered burst-pause responses in adult animals exposed to alcohol (Li et al., 2025; Ma et al., 2022) and suggest CINs may show altered responses to salient stimuli in PAE mice. These results highlight DMS CIN dysfunction as particularly relevant to behavioral impairments observed in FASD. Complementing our electrophysiological findings, we observed significantly reduced ACh release within the DMS in PAE offspring. This reduced release of ACh may be due to both the reduction of DMS CIN numbers and lowered CIN action potential rate, as CINs are the principal source of ACh in the striatum.

Our analyses of operant behavior show that PAE mice exhibit diminished behavioral flexibility, a behavior mediated in part by DMS cholinergic circuitry (Bradfield et al., 2013; Ma et al., 2022; Ragozzino et al., 2009). While initial instrumental A-O contingency learning was unaffected by PAE, adaptation to new contingencies during reversal learning was disrupted in PAE offspring. This selective impairment in reversal learning is in agreeance with previous rodent studies showing reversal learning deficits in T-Maze (Lee and Rabe, 1999; Thomas et al., 2004) and operant discrimination responding (Marquardt et al., 2014) following developmental alcohol exposure. Additionally, our observed increase in compulsive alcohol consumption in PAE offspring aligns with clinical studies showing that PAE in humans is a risk factor for developing symptoms of AUD (Baer et al., 1998; Goldschmidt et al., 2019), which is characterized by compulsive drinking despite negative consequences (Koob and Volkow, 2010). This compulsivity may also be an aspect of result from broader deficits in inhibitory control and decision-making, which is known to be disrupted by PAE (Fryer et al., 2007; Olguin et al., 2021). Such inflexibility is a hallmark of FASD pathology in humans (Green et al., 2009; Mattson et al., 1999; McGee et al., 2008), and our findings implicate DMS cholinergic dysfunction in potentially mediating this critical deficit.

Given the essential role of ACh in modulating MSN activity and mediating behavioral flexibility (Gangal et al., 2023; Huang et al., 2024; Ma et al., 2022), decreased cholinergic signaling likely contributes to the impaired cognitive functions seen in this study. Indeed, reduced cholinergic tone in the DMS has been implicated in diminished flexibility and compromised decision-making processes (Bradfield et al., 2013; Huang et al., 2024; Ma et al., 2022; Ragozzino et al., 2002), providing a possible mechanistic link between the observed cellular deficits and behavioral impairments. In particular, our finding that the pause response following burst excitation in CINs is shortened by PAE is especially relevant. This finding is consistent with previous work implicating shortened CIN pause responses after adult alcohol exposure with impaired behavioral flexibility (Li et al., 2025; Ma et al., 2022). CIN burst-pause responses are driven by changes in contingency and salient stimuli, and pauses in CIN activity are thought to allow for new corticostriatal plasticity to occur (Reynolds et al., 2022; Zucca et al., 2018). Thus, the attenuated pause response, along with an overall reduction in ACh tone, may be responsible for the reversal learning deficits observed in PAE mice. The altered CIN pause response may similarly explain the inflexible drinking seen in PAE offspring, as the salience of quinine is likely to elicit burst and pause firing in CINs, which are known to modulate alcohol drinking (Sharma et al., 2024). Furthermore, the reduction in spontaneous ACh release may also contribute to this phenomena, as DMS MSNs are strongly modulated by ACh and are integral for alcohol drinking behavior (Cheng et al., 2017). Lastly, our results are broadly supportive of multiple studies showing improved cognition following PAE when choline, a precursor to ACh, supplementation is taken during development (Ernst et al., 2022).

In conclusion, this study identified a novel and relevant deficit in DMS cholinergic circuitry and identified associated cognitive and behavioral impairments in adult PAE offspring. The specific reduction in CIN populations and functional impairments in cholinergic signaling highlight striatal cholinergic neurons as a promising target for therapeutic intervention. Future research aimed at restoring cholinergic function or enhancing CIN activity could yield novel approaches to mitigating cognitive deficits and reducing compulsive behaviors associated with FASD.

## Acknowledgements

This research was supported by NIAAA R01AA021505 (JW), U01AA025932 (JW) R01AA027768 (JW), R01AA030293 (JW), and R01AA024659 (RM). We thank Dr. Siara Rouzer for assisting with the vapor alcohol exposure procedure and Dr. Amanda Essoh for assisting with live brain slice preparation.

## Financial Disclosures

All authors report no biomedical financial interests or potential conflicts of interest.

## Notes

### Competing Interest Statement

The authors have declared no competing interest.

